# Super Resolved Single-Cell Spatial Metabolomics from Multimodal Mass Spectrometry Imaging guided by Imaging Mass Cytometry

**DOI:** 10.1101/2024.10.21.619323

**Authors:** Efe Ozturk, Abhijeet Venkataraman, Felix G. Rivera Moctezuma, Ahmet F. Coskun

## Abstract

Mass spectrometry imaging (MSI) is a powerful technique for spatially resolved analysis of metabolites and other biomolecules within biological tissues. However, the inherent low spatial resolution of MSI often limits its ability to provide detailed cellular-level information. To address this limitation, we propose a guided super-resolution (GSR) approach that leverages high-resolution Imaging Mass Cytometry (IMC) images to enhance the spatial resolution of low-resolution MSI data. By using these detailed IMC images as guides, we improve the resolution of MSI images, creating high-resolution metabolite maps. This enhancement facilitates more precise analysis of cellular structures and tissue architectures, providing deeper insights into super-resolved spatial metabolomics at the single-cell level.

## Introduction

MSI has emerged as a crucial tool in spatial metabolomics, allowing for detailed mapping of chemical compounds across biological tissues^1,2^. MSI techniques, such as Matrix-Assisted Laser Desorption/Ionization (MALDI)^3,4^ or Time-of-Flight Secondary Ion Mass Spectrometry (TOF-SIMS)^5^ offer the ability to simultaneously detect and visualize a wide range of metabolites, lipids, and proteins within tissue sections. Despite these advantages, a significant limitation of MSI is its relatively low spatial resolution compared to other imaging modalities like IMC, due to the diffraction limit and the size of the laser or ion beam used in MSI^6,7^. This inherently limits the spatial resolution, often making it difficult to capture fine cellular details or resolve individual cells within tissue sections. In contrast, IMC achieves higher resolution by detecting metal-tagged antibodies in a much more localized manner, enabling more precise single-cell analysis and detailed tissue architecture studies^8,9^.

IMC offers images at high resolution, capturing specific markers and providing further insight into tissue biochemistry. By integrating the high-resolution information from IMC with the rich chemical data from MSI, we can overcome the spatial resolution constraints of MSI and achieve a more comprehensive understanding of tissue biochemistry.

Several computational methods have been developed to improve the spatial resolution of MSI, both in single-modal and multimodal settings. Single-modal approaches often rely on methods that are developed using MSI data alone to enhance image resolution^10–12^. While these techniques can provide some improvement, they are often limited by the inherent resolution constraints of MSI and may fail to capture the detailed structures needed for single-cell or subcellular analysis.

To address these limitations, multimodal super-resolution approaches have emerged, where high-resolution images from complementary modalities serve as guides to enhance low-resolution MSI data^13–15^. In these methods, the high-resolution images provide structural or phenotypic context, which guides the reconstruction of super-resolved MSI images, resulting in more accurate metabolite maps that preserve both spatial and chemical integrity.

In this work, we introduce a GSR method to enhance the spatial resolution of low-resolution MSI images using high-resolution IMC images. Several GSR methods have been developed in computer vision for enhancing image resolution across different applications^16–20^. Our method employs an atention mechanism^21^ to selectively emphasize the most relevant features in the IMC guide images, improving the accuracy of the super-resolved metabolite maps. This GSR approach enables the creation of high-resolution metabolite maps that retain both the chemical specificity of MSI and the spatial fidelity of IMC. This method not only improves the visualization of metabolites at the cellular level but also enhances the overall interpretability of MSI data for beter analysis of both cellular and larger tissue structures.

## Results

### IMC-guided Super Resolution MALDI

The analysis of the colorectal cancer dataset^22^ demonstrates that the GSR approach effectively enhances the resolution of low-resolution MSI data by leveraging high-resolution IMC as a guide. The acquisition of both low-resolution MSI and high-resolution IMC data provides the foundation for this process (Fig. 1-a). After selecting the metabolites with significant spatial information, which are chosen automatically (Fig. 1-b), based on entropy and nonzero-pixel to zero-pixel ratio thresholds (Methods: Automated Metabolite Selection), the GSR method produces super-resolved metabolite maps that reveal finer spatial details compared to the original low-resolution MSI (Fig. 1-c, Methods: Guided Super Resolution). These enhanced maps allow for beter visualization of metabolite distribution and tissue architecture at the cellular level. In particular, the super-resolution outputs highlight sharp boundaries and improved spatial fidelity, especially in complex tissue regions (Fig. 1-d).

**Figure 1.**
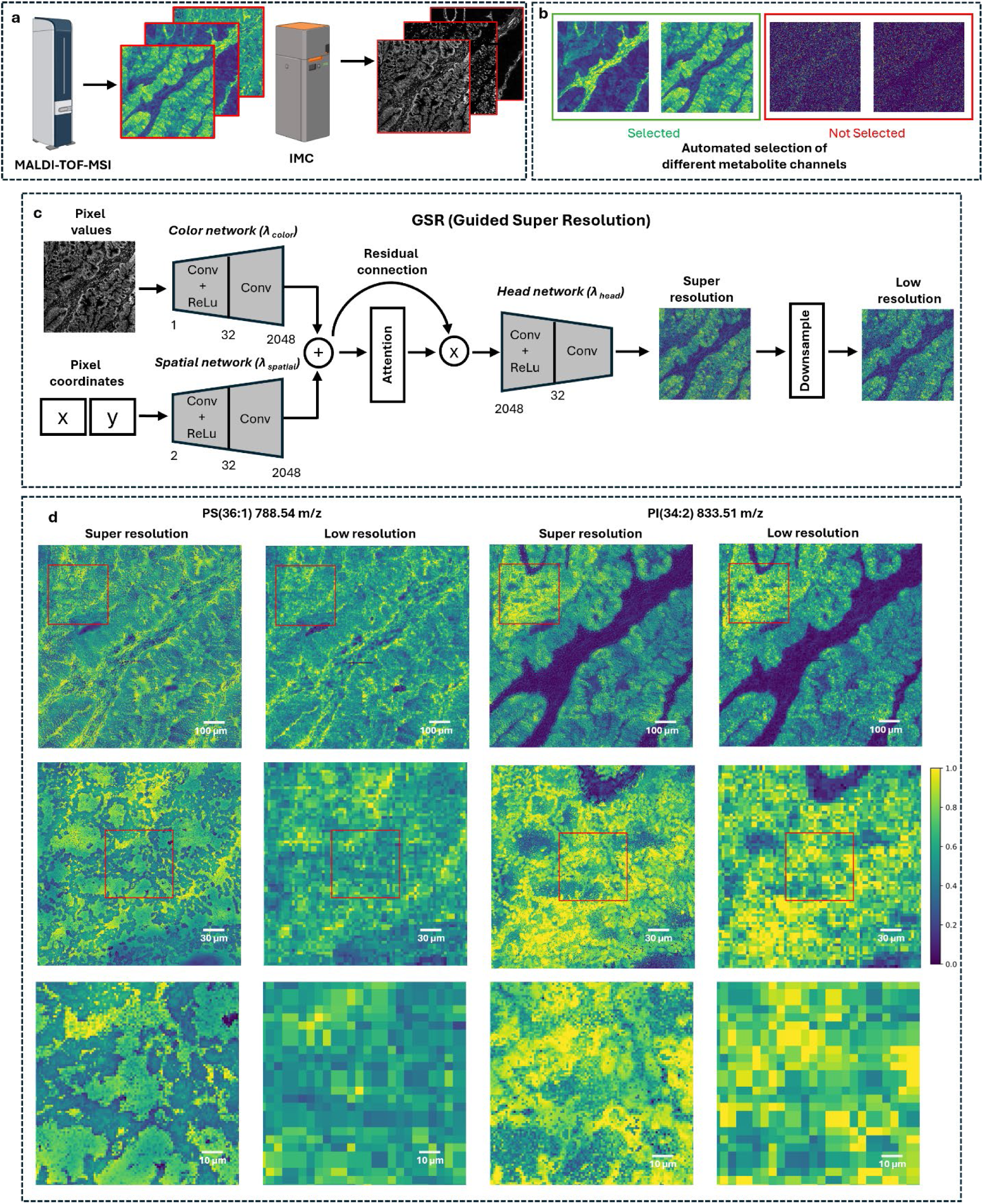
Workflow for GSR in high-resolution IMC and MALDI MSI for colorectal cancer analysis. (a) Acquisition of low-resolution MSI data via MALDI and high-resolution IMC images. (b) Automated selection of metabolite channels from MSI data, with selected channels (green) showing significant information and non-selected channels (red) being excluded. (c) The GSR process integrates spatial coordinates and pixel intensities from the IMC images and produces super-resolution MSI images. Downsampling of the super-resolution output is used to calculate loss by comparing it with the original low-resolution MSI. (d) Examples of two GSR outputs are phosphatidylserine (PS 36:1, 788.54 m/z) and phosphatidylinositol (PI 34:2, 833.51 m/z). The top row shows super-resolution and low-resolution images for larger regions, while the bottom row presents the corresponding zoomed-in regions of interest.

Phosphatidylserine (PS 36:1) and phosphatidylinositol (PI 34:2) have critical roles in cell membrane composition and signaling, respectively. Phosphatidylserine is one of the most abundant anionic phospholipids in eukaryotic cell membranes, accounting for roughly 10% of the total lipid composition^23^. This lipid plays an essential role in maintaining the structural integrity of the cell membrane. On the other hand, phosphatidylinositol is a key signaling molecule involved in glucose metabolism and insulin signaling, which are vital for maintaining the cell’s energy balance^24^. The ability to super-resolve these metabolite distributions offers greater insight into their spatial localization and potential roles in cancer cell metabolism and signaling pathways. The comparison between GSR outputs and low-resolution data confirms that the GSR method preserves the chemical specificity of metabolites while providing high-resolution detail, which is critical for detailed biological analysis at the cellular level.

The results demonstrate the effectiveness of the GSR approach in significantly improving the spatial resolution of MSI (Fig. 2). The GSR method enhances the clarity of cellular structures in the selected ROI, revealing sharper details and more defined boundaries compared to the low-resolution MSI (Fig. 2a, b). This improvement is quantitatively confirmed by the line profile comparison, where the super-resolved image shows two distinct peaks for two single cells with FWHM values of 4 pixels each, while the low-resolution image presents a single, broader peak with an FWHM of 43 pixels (Fig. 2c). This represents an approximately five-fold increase in resolution (5.375), demonstrating that the super-resolved image captures individual cell details at a significantly higher resolution.

**Figure 2.**
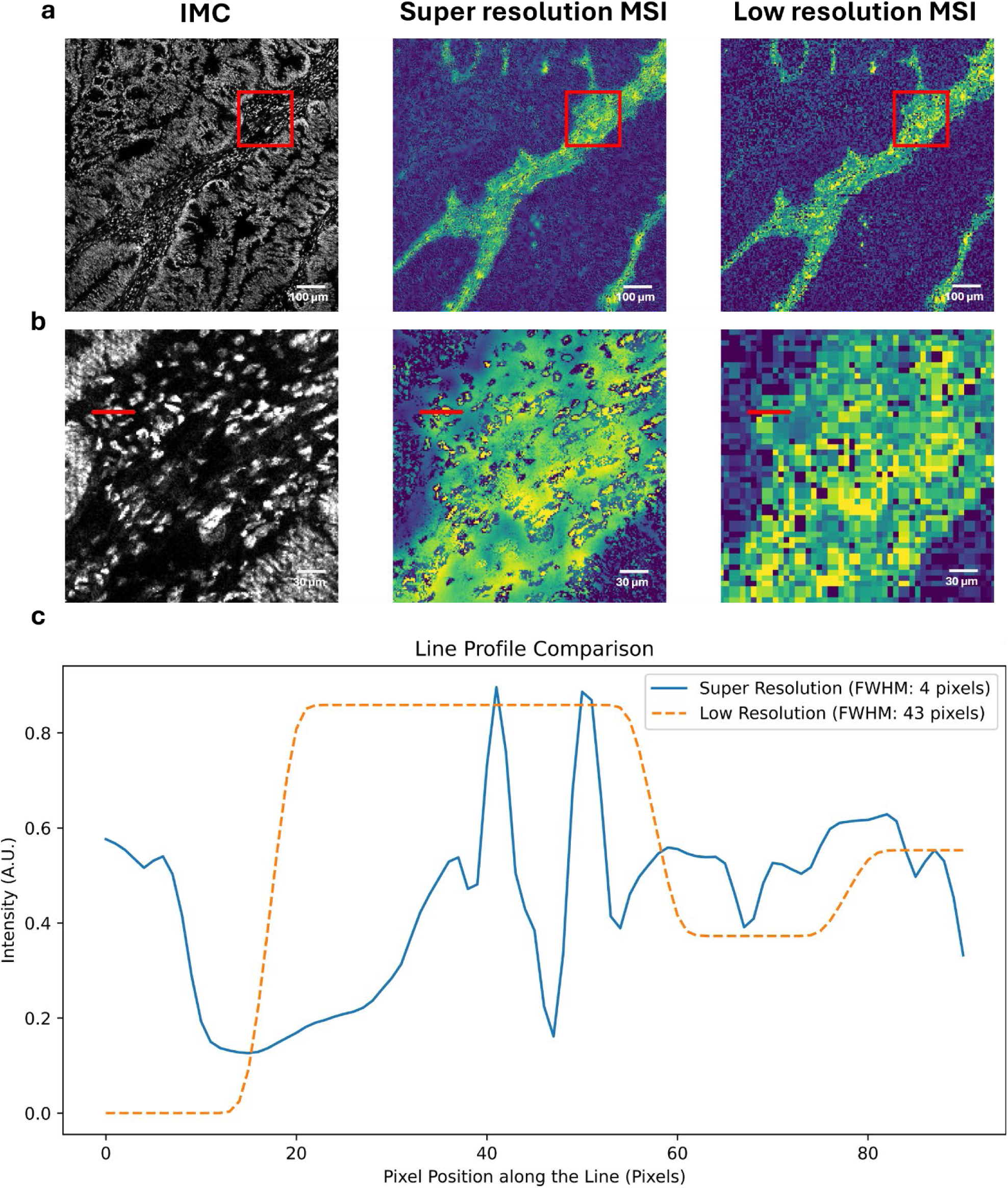
Comparison of low-resolution and super-resolution MSI data. (a) The first panel (left) displays the high-resolution IMC image, while the middle and right panels show the super-resolved and original low-resolution MSI images of PI(38:4), respectively. The red bounding box highlights the region of interest (ROI) used for further analysis. (b) Zoomed-in view of the ROI from (a), with a red line drawn over a cell in both the IMC and MSI images, indicating the region along which the intensity profiles are compared. (c) Line profile comparison of the intensity values along the red line from the low-resolution and super-resolved MSI images. The solid blue line represents the super-resolved image, while the dashed orange line shows the low-resolution MSI. The full width at half maximum (FWHM) is calculated for both profiles.

The clustering of cells based on metabolite expression reveals distinct metabolic profiles across different cell populations (Fig. 3-a). This clustering is further highlighted by the heatmap, which underscores key metabolites that drive the separation of these populations (Fig. 3-b). When overlaid with cell phenotypes, the analysis reveals strong correlations between certain cell types and their metabolic states (Fig. 3-c). For this analysis, we used the DNA IMC channel to guide the GSR outputs for each metabolite. Then, the original cell mask, validated for consistency, was applied to obtain the cell contours and compute the mean intensity for each cell. This approach ensures accuracy in cell identification and allows for meaningful comparison across spatially resolved metabolic profiles. The spatial distribution of these phenotypes in the tissue further supports this, showing how different cell types organize within the tissue architecture (Fig. 3-d). This analysis indicates that the GSR-enhanced metabolite maps provide a meaningful resolution that captures and differentiates metabolic variability at the cellular level, offering deeper insights into tissue heterogeneity. The mean intensity plots for selected metabolites also reinforce the metabolic distinctions across different cells (Fig. 3-e).

**Figure 3.**
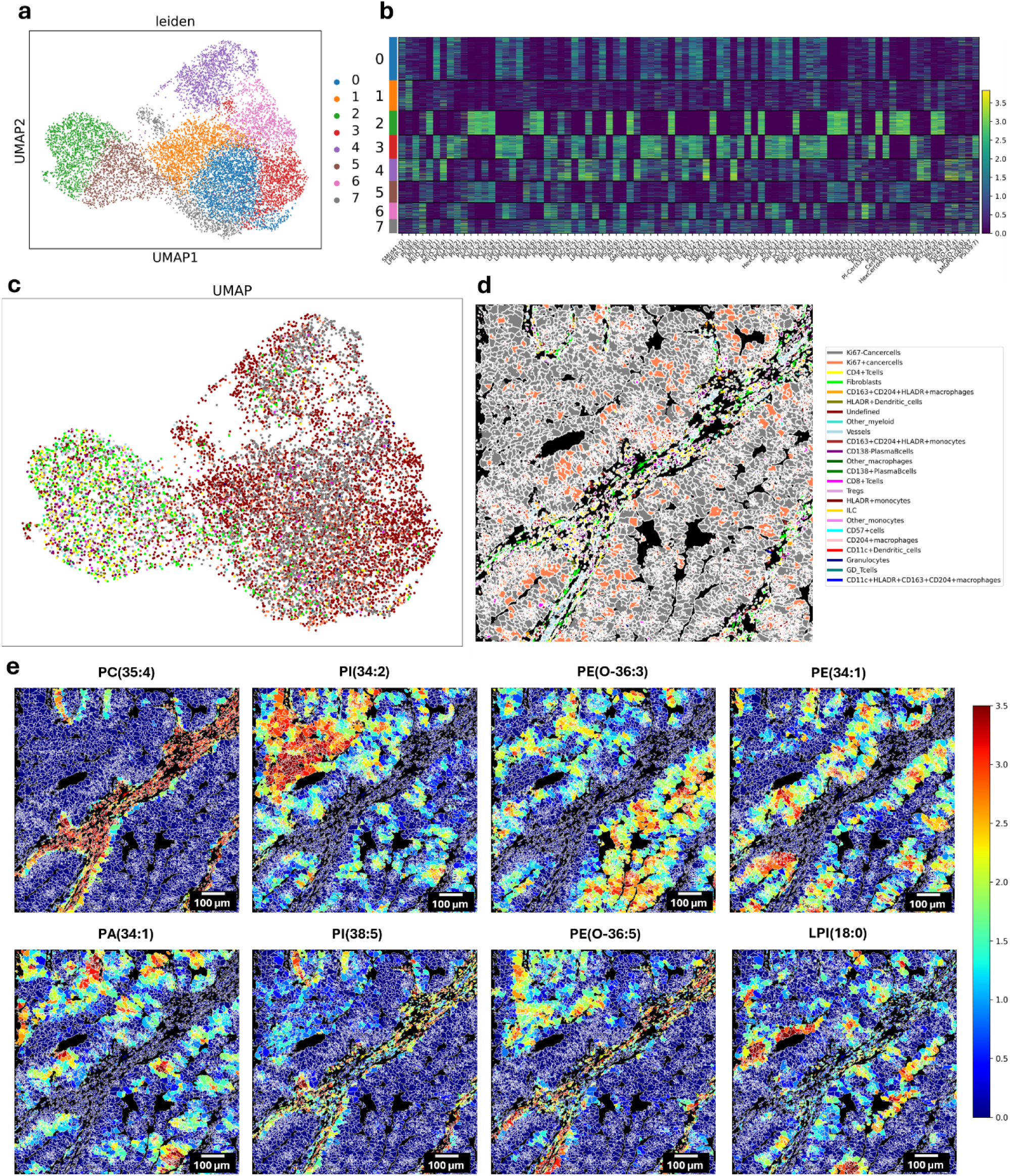
Clustering of Cells Based on Metabolites and Visualization of Cell Phenotypes in UMAP and Tissue Images. (a) UMAP plot showing the clustering of cells based on selected metabolite intensities, colored by Leiden clustering labels. (b) Heatmap displaying the expression of selected metabolites across the different clusters. (c) UMAP plot colored based on cell phenotypes, showing how different cell types are distributed across the metabolite-based clusters. (d) Spatial visualization of the same cell phenotypes overlaid on the tissue image, highlighting the distribution of different cell types within the tissue. (e) Mean intensity plots show the distribution of metabolites across the cells, with each subplot representing a different metabolite intensity. The cell mask and the corresponding phenotypes used for clustering and visualization were derived from the original data, ensuring consistency in cell identification and spatial analysis.

The selection of different IMC channels as guides impacts the final super-resolved MSI output (Fig. 4). The ability of each guide to emphasize certain spatial features within the mass spectrometry images is evident, as shown in the varying levels of contrast and clarity across rows (Fig. 4-a to Fig. 4-e). Some guides highlight cellular boundaries more distinctly (e.g., DNA in Fig. 4-a), while others bring out subtle intra-cellular variations (e.g., Beta-catenin in Fig. 4-c), suggesting that selecting appropriate guides can be tailored to specific analysis goals—whether for cell segmentation, tissue architecture, or intracellular feature detection. Notably, in the zoomed-in regions (third and fourth columns), the GSR approach demonstrates its capability to resolve fine details that were previously obscured in the low-resolution MSI. The sharpness of cellular boundaries and the emergence of distinct structures in high-resolution regions emphasize the strength of the GSR method in single-cell metabolomics. The variations observed between different guides suggest that incorporating multiple guide channels could provide a more holistic view of the spatial metabolite distribution, capturing both large-scale tissue organization and fine cellular details. Moreover, the differences across rows highlight the versatility of the GSR framework. Depending on the biological question—whether one is interested in identifying specific cellular structures or broader tissue regions—selecting the optimal IMC guide could significantly enhance the interpretability of the data.

**Figure 4.**
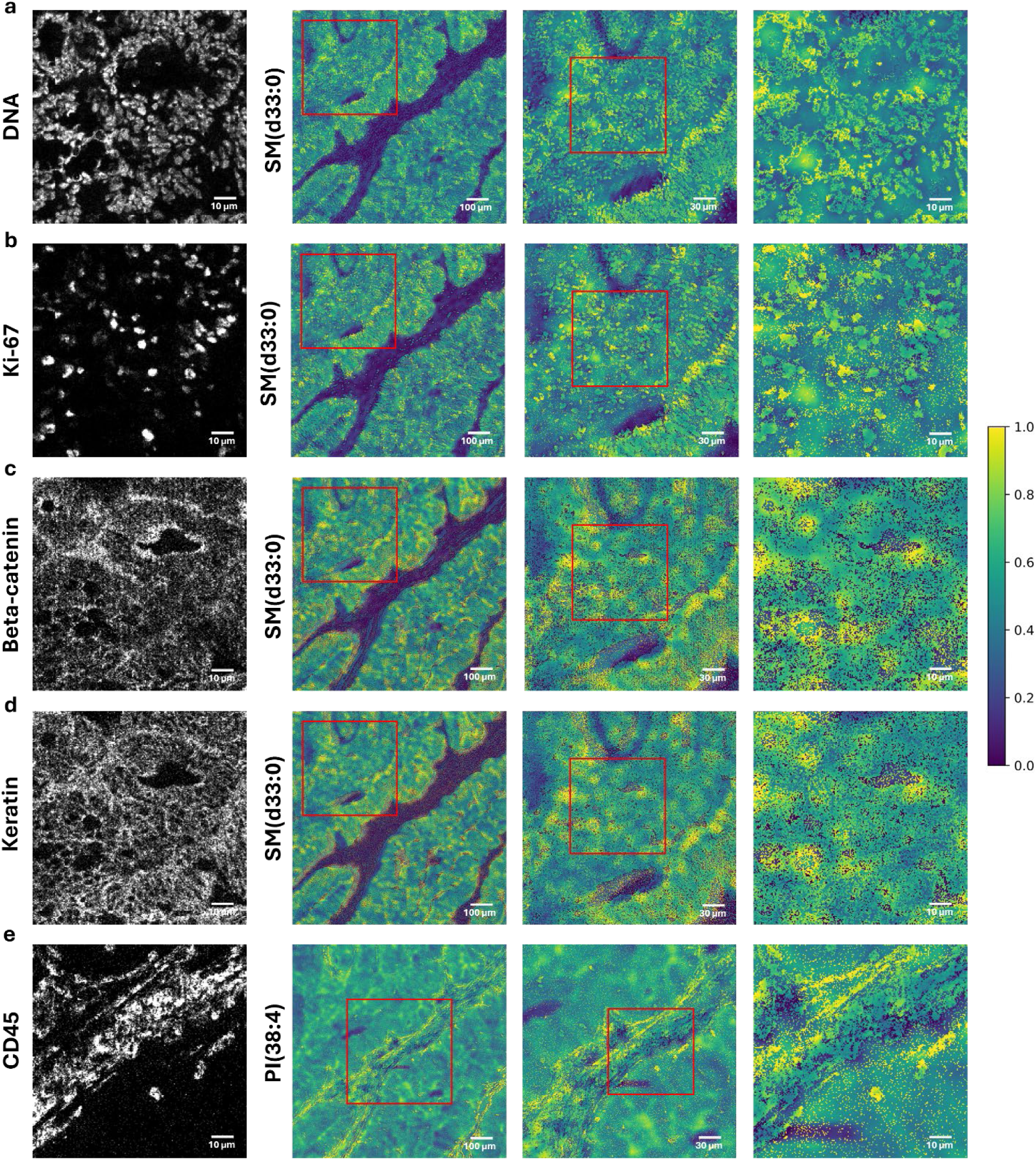
GSR results demonstrating the impact of using different IMC channels as guides. Each row corresponds to a distinct IMC guide: (a) DNA, (b) Ki-67, (c) Beta-catenin, (d) Keratin, and (e) CD45. The first column displays grayscale IMC images of these markers. The second column shows the larger region of the resulting super-resolved mass spectrometry images, the third column provides a zoomed-in view of a region of interest, and the fourth column presents a further zoomed-in view within the zoomed region. The color bar on the right represents normalized metabolite intensity values. For DNA, Ki-67, Beta-catenin, and Keratin, SM(d33:0) is used as the metabolite. For CD45, PI(38:4) is used.

The comparison between cell masks generated from different sources (Fig. 5-a to Fig. 5-c). and their respective cell mean intensity maps (Fig. 5-d) provides insight into the accuracy and reliability of segmentation approaches. The first column shows the mask derived from the original dataset, serving as a baseline for comparison. The second column uses high-resolution IMC images to generate the mask, integrating nuclear (Ki-67) and cytosolic (vimentin and keratin) markers, yielding detailed cellular boundaries and clear nuclear-cytoplasmic differentiation. The third column presents an innovative approach where the guided super-resolution (GSR) outputs of the same markers (Ki-67, vimentin, and keratin) are used to create the mask. Notably, the GSR-based mask reveals more cells than the original mask from the baseline data, especially in regions where smaller or less distinct cells were missed. This suggests that the GSR method not only enhances the spatial resolution but also increases the level of detail captured, enabling the discovery of additional cells that may have been overlooked in the original low-resolution segmentation. The increase in detected cells highlights the potential of GSR to uncover previously hidden biological structures, improving the scope of single cell analysis. The cell mean intensity maps correspond to the cell masks in the third row, further validating the GSR method, as the additional cells detected by the GSR-based mask display coherent and biologically relevant intensity paterns. The close resemblance of the intensity distributions between the GSR and IMC-based maps emphasizes the reliability of GSR for accurate segmentation and improved cellular resolution. This enhancement is crucial for single-cell spatial metabolomics, where detecting and analyzing finer structural details can lead to deeper insights into tissue heterogeneity and the discovery of new cellular subtypes.

**Figure 5.**
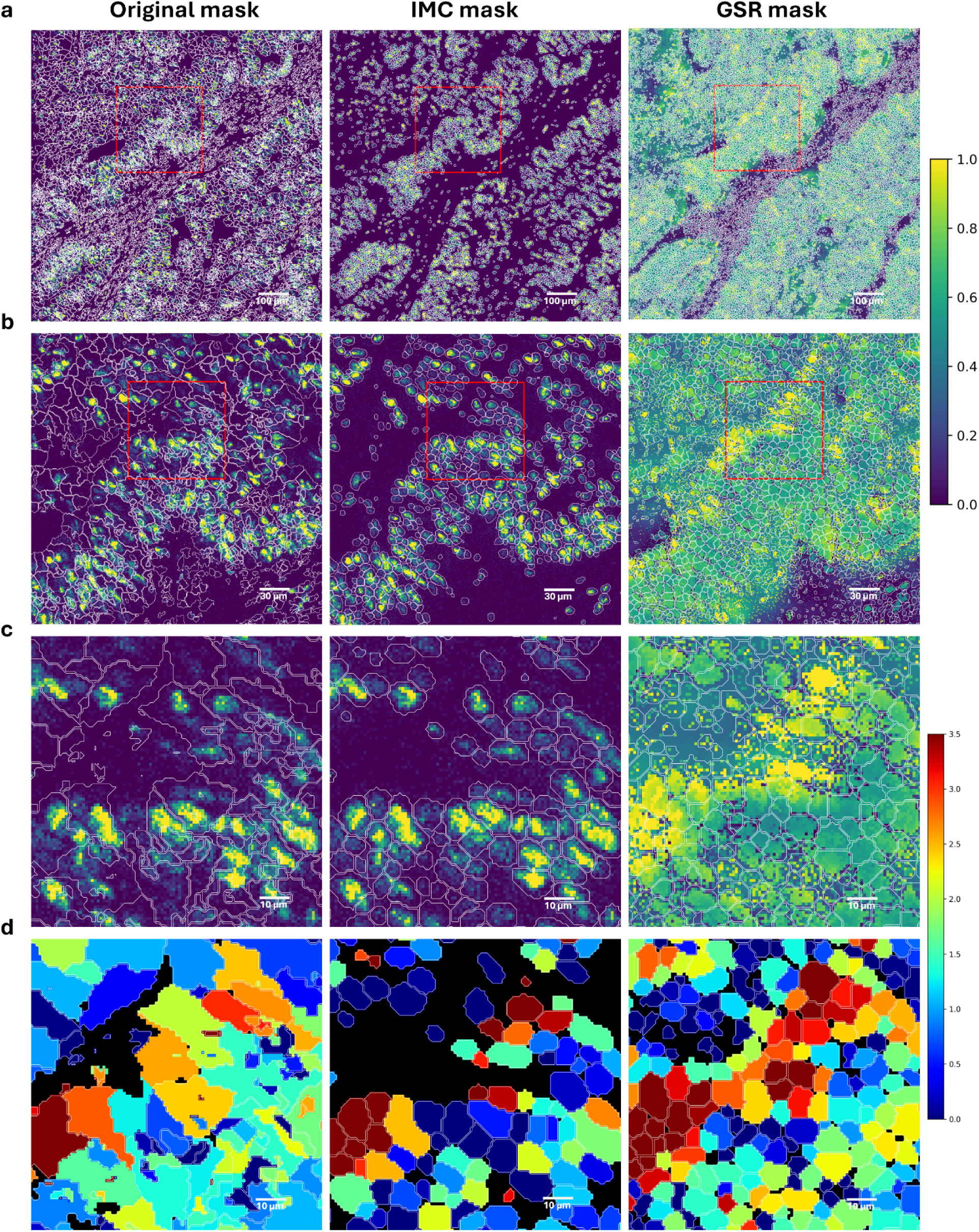
Comparison of segmentation masks and cell mean intensity maps across different regions and methods. (a) The larger tissue area is segmented using three different approaches. The first column presents the segmentation mask from the original dataset, the second column shows segmentation using IMC channels (Ki-67 for nuclei, vimentin, and keratin for cytosolic markers), and the third column displays segmentation based on GSR outputs, replacing IMC data. (b) The zoomed-in region from the larger tissue area is segmented with the methods used in (a). (c) Further zoomed-in region from the smaller tissue area segmented using the same approaches. (d) The corresponding cell mean intensity maps for the segmentation results from (c), highlighting the differences between the original dataset, IMC-based segmentation, and GSR-based segmentation. The metabolite used for this analysis is SM(d33:0).

The GSR outputs and corresponding cell mean intensity maps for two metabolites, PS(36:1) and PI(38:4), show that the segmentation method can be applied to various tissues while retaining critical structural details (Fig. 6). The consistency of the cell boundaries across all samples emphasizes the robustness of the GSR-based approach. The mean intensity maps offer insights into how metabolite distributions differ between cells and tissues, highlighting the variability in intensity levels across tumor samples. GSR-based segmentation reveals subtle intra-cellular variations that could correspond to different metabolic states, further supporting its potential to uncover nuanced tissue heterogeneity. This closer view into cell-level metabolic paterns provides deeper insights into tumor-specific metabolomic profiles. The ability to apply the same GSR-based cell mask consistently across different tumor samples (Fig. 6-a to Fig. 6-e) reinforces the generalizability of this approach for spatial metabolomics. This flexibility is crucial for ensuring reproducibility across studies involving varied biological samples, offering a tool for comparative analysis in clinical and research settings.

**Figure 6.**
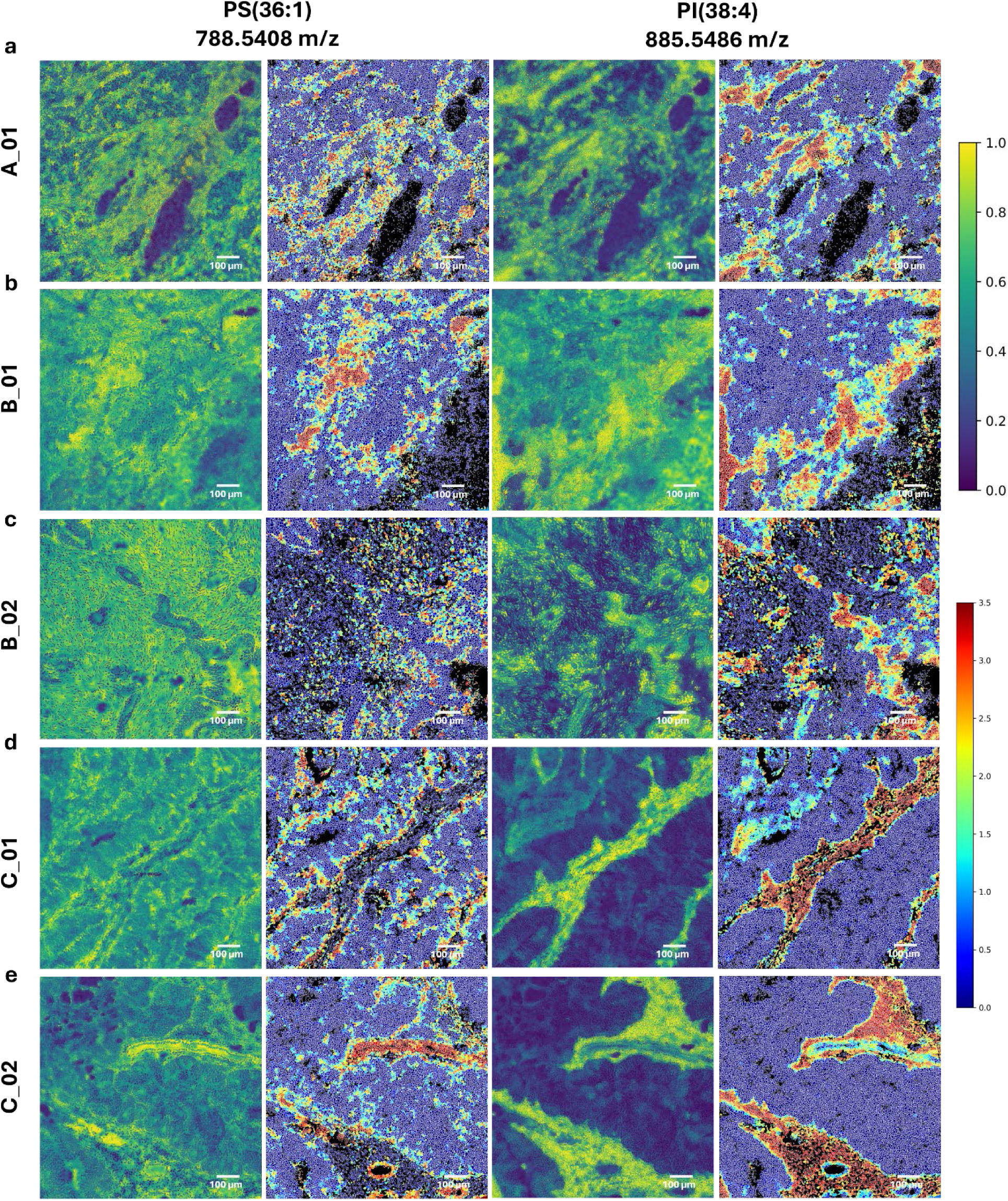
Comparison of GSR outputs and corresponding cell mean intensity maps across different tumor samples. (a), (b), (c), (d), and (e) correspond to tumor samples A_01, B_01, B_02, C_01, C_02, respectively. The first and second columns display the super-resolved GSR output and the corresponding cell intensity map for PS(36:1), while the third and fourth columns show the super-resolved GSR output and cell intensity map for PI(38:4). The color bar on the right represents the normalized intensity values. Cell masks were generated using GSR output of DNA as the nuclei channel and GSR outputs of vimentin + keratin for the cytosolic regions.

The combined visualization of UMAPs and metabolite intensity maps reveals clear tissue-specific metabolic signatures associated with distinct cell phenotypes (Fig. 7). The UMAPs, which focus on each tissue (Fig. 7-a to Fig. 7-e), show that metabolic heterogeneity aligns with specific immune cell populations, particularly within macrophages and monocytes. This suggests that these cell types play a significant role in shaping the metabolic landscape within each tissue. The intensity maps on the left indicate notable variations in metabolite levels across tissues, reflecting how local tissue environments influence metabolic activity. Meanwhile, the heatmaps provide further insight into how these metabolic changes correlate with the presence of specific cell types, with some macrophage and monocyte subtypes showing elevated levels. These observations highlight the capacity of GSR to capture cellular metabolic differences.

**Figure 7.**
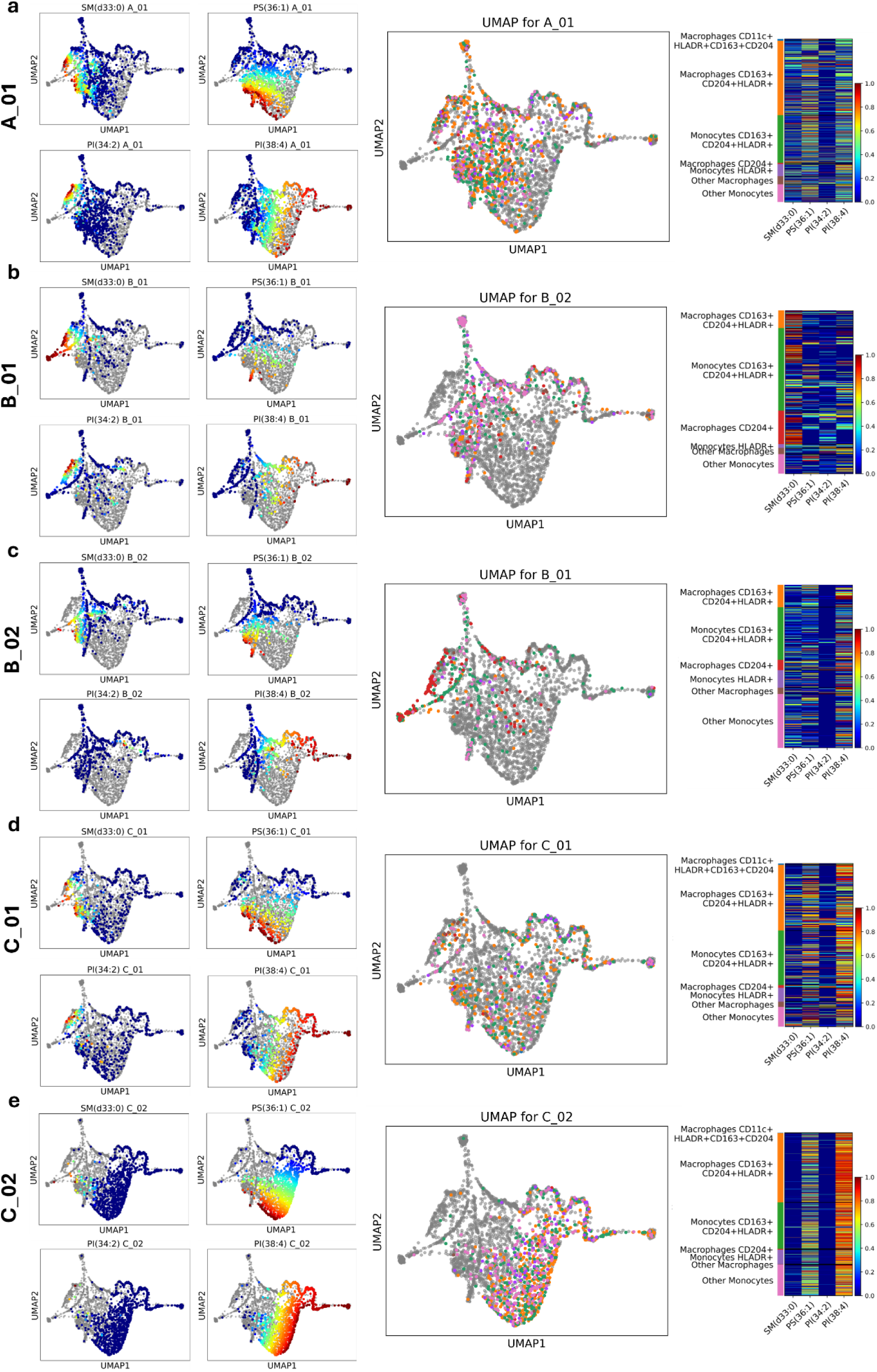
Combined UMAPs and metabolite intensity maps across different tissue samples. (a), (b), (c), (d), and (e) correspond to tumor samples A_01, B_01, B_02, C_01, C_02, respectively. The left panels display intensity maps for four selected metabolites (SM(d33:0), PS(36:1), PI(34:2), and PI(38:4)) for each tissue sample. In the middle panels, UMAPs are colored by phenotypes, with cells from other tissues grayed out to emphasize the tissue-specific phenotypic distribution. The right panels show corresponding heatmaps, presenting the relative intensity of the four metabolites across the selected phenotypes.

### IMC-guided Super Resolution TOF-SIMS

To evaluate the GSR method, we used TOF-SIMS data from human lung cancer tissue^25^, which includes sufficiently high-resolution MSI (Fig. 8). Because of the high-resolution MSI images, we used it as a benchmark to evaluate the performance of the GSR method quantitatively. We downsampled the original high-resolution MSI images to create low-resolution versions, trained the algorithm using these low-resolution images with IMC guides, and generated super-resolved outputs, which were compared to the original high-resolution MSI data (Fig. 8a). The super-pixel maps provide a pixel-wise comparison across low-resolution, original high-resolution, and super-resolved data. Quantitative analysis comparing pixel intensities between the super-resolution and original high-resolution images shows that the GSR approach outperforms baseline interpolation techniques like bilinear, bicubic, and nearest neighbors, with superior PSNR, SSIM, and MSE scores (Fig. 8b). The line profile comparison further demonstrates that GSR accurately reproduces the intensity paterns seen in the high-resolution image (Fig. 8c). These results confirm that GSR is highly effective at preserving both the spatial and chemical integrity of the data, making it a reliable method for enhancing MSI resolution and improving downstream analyses.

**Figure 8.**
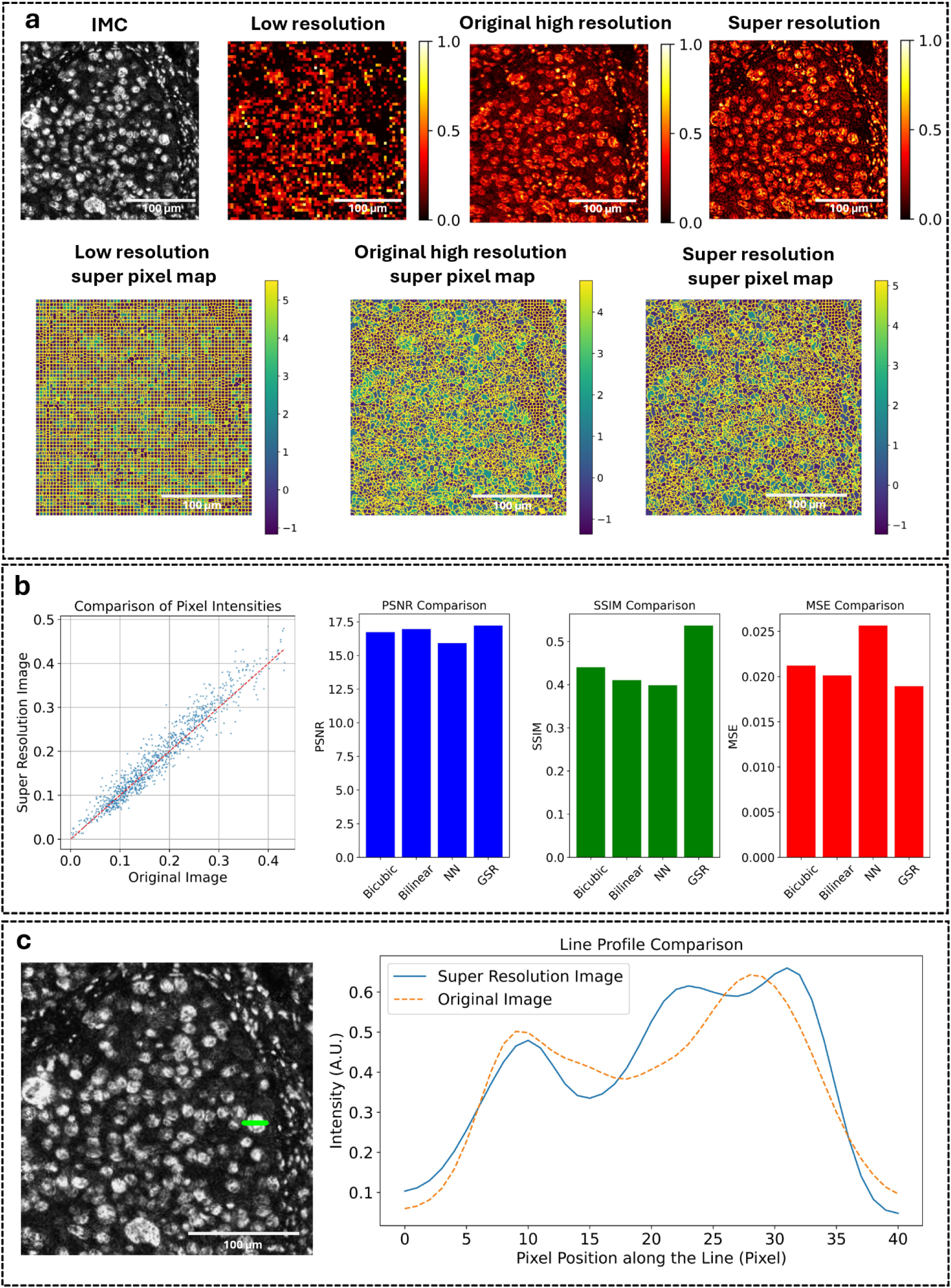
Comparison of super-resolution with ground truth high-resolution MSI in human lung cancer. (a) Visual comparison between low-resolution, original high-resolution, and super-resolution TOF-SIMS data. The IMC image is used as a guide for super-resolution. Below, super-pixel maps are shown for low-resolution, original high-resolution, and super-resolution data, providing a pixel-wise comparison. (b) Quantitative comparison of pixel intensities between the super-resolution image and the original high-resolution image. Scatter plots and evaluation metrics (PSNR, SSIM, and MSE) are used to compare the performance of the proposed GSR method against baseline techniques, including bicubic, bilinear, and nearest neighbors (NN). (c) Line profile comparison between the super-resolution image and the original high-resolution image along a selected region (highlighted as green) demonstrates the correspondence of intensity patterns between the two.

The GSR method effectively enhances the spatial resolution of TOF-SIMS metabolite maps across four additional human lung cancer tissue regions, as demonstrated by the reconstructed outputs (Fig. 9). In each region, the super-resolved images (Fig. 9-d) closely match the original high-resolution TOF-SIMS data (Fig. 9-c), successfully capturing fine cellular structures and tissue architecture that are blurred in the low-resolution images (Fig. 9-b). The high-resolution IMC images (Fig. 9-a) were used as guides for this process. The pixel intensity comparison plots (Fig. 9-e) provide further validation, showing a strong correspondence between the super-resolved and original high-resolution images, and highlighting the precision of the GSR method in preserving spatial information.

**Figure 9.**
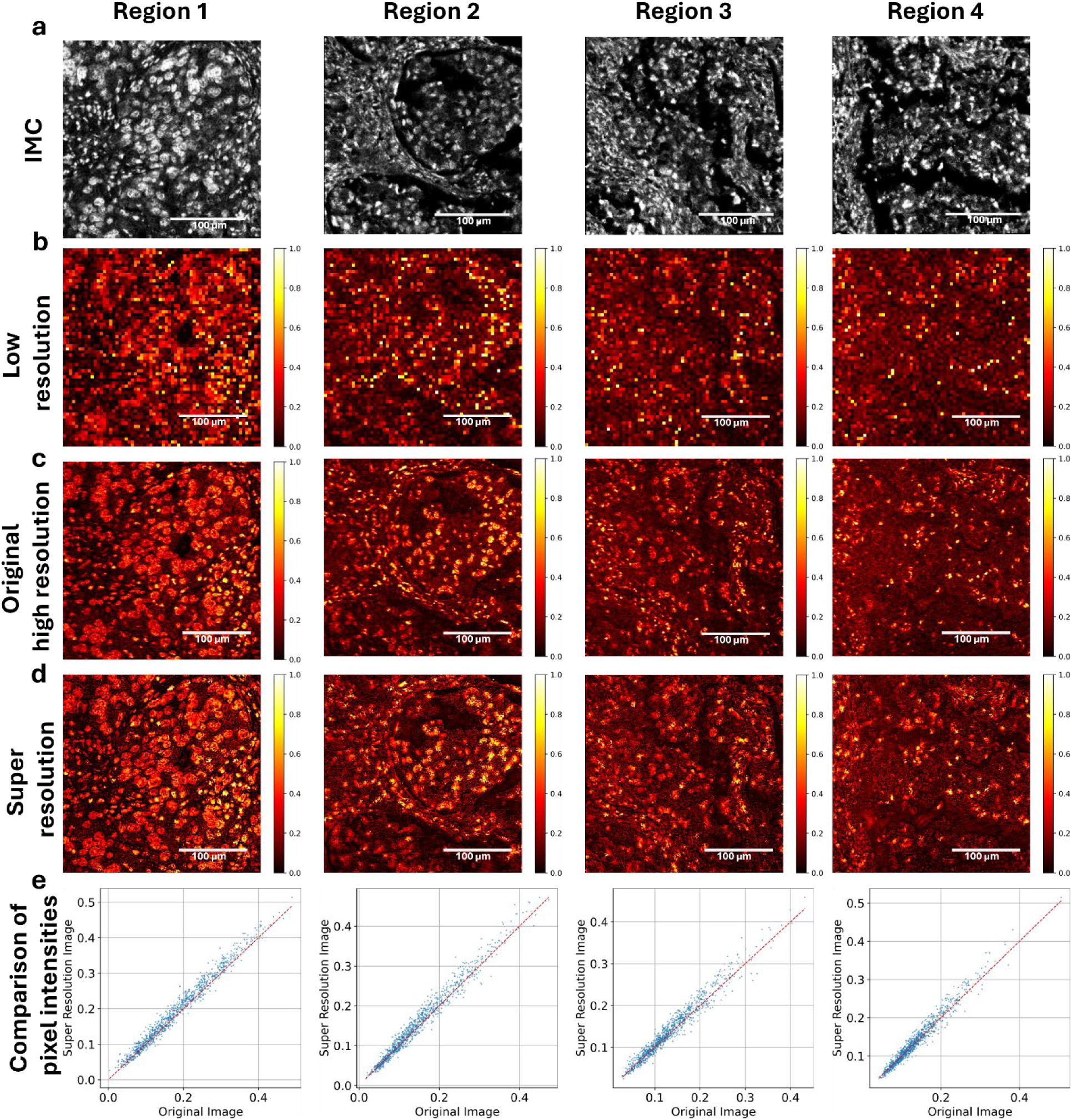
Comparison of IMC, low-resolution, original high-resolution, and GSR super-resolution TOF-SIMS images across four distinct human lung cancer tissue regions. (a) High-resolution IMC images used as guides. (b) The corresponding low-resolution TOF-SIMS images. (c) The original high-resolution TOF-SIMS images were used as ground truth for comparison. (d) The GSR super-resolved TOF-SIMS images demonstrate an enhanced spatial resolution that closely matches the original high-resolution images. (e) Pixel intensity comparison plots, with each scatter plot comparing pixel intensities between the super-resolved and original high-resolution images.

## Methodology

### Guided Super Resolution

To achieve the guided super-resolution of low-resolution MSI images, we enhance low-resolution MSI data using high-resolution guide images from IMC. The low-resolution MSI image, which contains spatially resolved metabolite data, serves as the source image and has dimensions *MxM*. The high-resolution IMC image, a grayscale image providing detailed structural information, acts as the guide and has dimensions *NxN* with *N* = *DxM* where *D* is the upsampling factor.

The pixel-to-pixel transformation involves two primary inputs for each pixel in the guide image: the grayscale pixel values for IMC images, along with spatial coordinates (x, y). Thus, the input to the network for each pixel is a vector [*Gray, x, Y*].

The network consists of two sub-networks that process the spatial and color inputs separately. The spatial network processes the spatial coordinates (x, y). It takes two input channels and first applies a 1*x*1 convolutional layer with 32 filters, followed by a ReLU activation. This is followed by a larger *kxk* convolutional layer with 2048 filters (with zero padding), where *k* is the kernel size, to capture spatial relationships over larger areas. This series of layers allows the network to extract meaningful spatial features from the input coordinates, retaining spatial consistency during the super-resolution process.

The color network takes the grayscale pixel values as the input. Similar to the spatial network, the color network applies a 1*x*1 convolutional layer with 32 filters, followed by a ReLU activation. The network then applies a *kxk* convolutional layer with 2048 filters, maintaining the same kernel size and padding as the spatial network.

After extracting the features from both the spatial and color networks, the outputs are merged. A channel atention mechanism is then applied to the merged features. This atention mechanism helps refine the merged outputs by dynamically weighing the most important channels. To achieve this, the atention mechanism first applies a 1*x*1 convolution that reduces the number of channels from 2048 to 512, followed by a ReLU activation. Another 1*x*1 convolution restores the channels back to 2048, and a sigmoid activation generates atention weights. These weights are multiplied with the merged features, ensuring that the most relevant information is emphasized before the final high-resolution MSI image is produced.

The merged and refined features are passed through a head network to generate the final high-resolution MSI output. The head network first applies a ReLU activation to the atention-refined features. This is followed by a *kxk* convolutional layer with 32 filters to further process the features and another ReLU activation. Finally, the network applies a 1*x*1 convolutional layer with 1 filter to produce the high-resolution MSI output. This final convolution ensures that the output is a single-channel high-resolution MSI image with dimensions *NxN*.

The training process of the model involves minimizing a reconstruction loss and applying regularization to the spatial, color, and head networks. The reconstruction loss ensures that the downsampled predicted high-resolution MSI image is consistent with the low-resolution source image. This loss is calculated using *L*_1_ loss as follows:

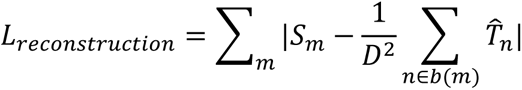

Where *b*(*m*) represents the block of *DxD* pixels in the high-resolution image 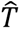 corresponding to the low-resolution pixel *S*_*m*_.

In addition to the reconstruction loss, the model incorporates regularization to prevent overfitting. Three separate regularization terms are applied to the spatial, color, and head networks. Each regularizer applies an *L*_2_ penalty on the weights of the respective network, controlling its complexity and ensuring that the learned mapping function is not overly complex. The regularization for the spatial, color, and head networks is defined as follows:

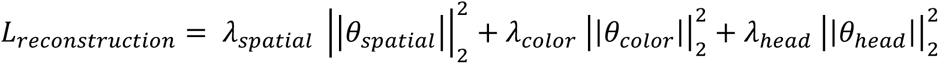

where *θ*_*spatial*_, *θ*_*color*_, and *θ*_*head*_ are the weights of the spatial, color, and head networks, respectively, and *λ*_*spatial*_, *λ*_*color*_, and *λ*_*head*_ are the regularization parameters.

The Adam optimizer is used to minimize this total loss during training. The training data consists of paired guide images (IMC) and source images (MSI). Once trained, the model is capable of predicting high-resolution MSI images, enabling the creation of detailed metabolite maps at single-cell resolution. Once trained, the super-resolution process involves using the network to predict the high-resolution MSI image. Each pixel in the guide image is passed through the model to generate the corresponding high-resolution MSI value, resulting in a super-resolved image 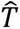 with dimensions *NxN*.

### Automated Metabolite Selection

To focus the GSR analysis on metabolites with significant spatial information, we applied a two-step selection process based on entropy and the nonzero-pixel to zero-pixel ratio. First, the entropy of each metabolite image was calculated to quantify the spatial complexity and variation of the metabolite distribution. Metabolites with higher entropy were considered to contain more meaningful spatial paterns. Second, we computed the ratio of nonzero-pixels to zero-pixels for each metabolite image to ensure that metabolites with widespread signals across the tissue, rather than sparse distributions, were selected. Adaptive thresholds for both metrics were established by calculating the mean minus the standard deviation across all metabolites in the dataset. Only metabolites with entropy and nonzero-pixel to zero-pixel ratio values above these thresholds were selected for further analysis. This approach allowed us to focus on metabolites with sufficient spatial variation, eliminating those with minimal or noisy signals.

### Segmentation Mask Generation

All segmentation masks used in this study were generated using the Mesmer model^26^. The whole-cell segmentation mode of Mesmer was applied, requiring one channel for nuclei and one channel for cytosol. Specifically, the Ki-67 IMC channel was used for nuclei segmentation, while vimentin and keratin channels were used for cytosolic segmentation, ensuring accurate delineation of whole cells.

## Conclusion

In this study, we introduced a GSR approach that leverages high-resolution IMC images to enhance the spatial resolution of low-resolution MSI data. Our results demonstrate that the GSR method significantly improves the visualization of metabolites at the single-cell level, providing a more detailed and precise representation of cellular structures and tissue architectures. By integrating spatial and pixel intensity information from IMC, the GSR algorithm successfully recovers high-resolution features while preserving the chemical specificity and spatial distribution of the metabolites.

Quantitative analysis confirmed that GSR outperforms traditional interpolation methods, such as bilinear and bicubic, in reconstructing high-resolution MSI data. Furthermore, GSR effectively replicates intensity profiles similar to those of ground truth high-resolution MSI, reinforcing its reliability and accuracy for spatial metabolomics.

The enhanced resolution provided by GSR enables more accurate clustering of cells based on metabolite expression, revealing distinct metabolic profiles that correspond to cell phenotypes. This capability holds great potential for uncovering new insights into tissue heterogeneity, cellular metabolic state and composition, and disease pathology, especially at the single-cell level.

Overall, GSR represents a powerful advancement in spatial metabolomics, allowing researchers to analyze MSI data with unprecedented clarity and resolution. Future work will focus on further optimizing the algorithm, exploring its applicability to other imaging modalities and tissue types, and expanding its use in diverse biological and clinical contexts to provide deeper insights into cellular biology and disease mechanisms.

## Acknowledgments

A.F.C. holds a Career Award at the Scientific Interface from Burroughs Wellcome Fund and a Bernie-Marcus Early-Career Professorship. A.F.C. was supported by start-up funds from the Georgia Institute of Technology and Emory University. This work was performed in part at the Materials Characterization Facility (MCF) at Georgia Tech. The MCF is jointly supported by the GT Institute for Materials (IMat) and the Institute for Electronics and Nanotechnology (IEN), which is a member of the National Nanotechnology Coordinated Infrastructure supported by the National Science Foundation (Grant ECCS-1542174). This work was performed in part at the Georgia Tech Institute for Electronics and Nanotechnology, a member of the National Nanotechnology Coordinated Infrastructure (NNCI), which is supported by the National Science Foundation (Grant ECCS-2025462). IEN also provided financial support in the form of a Facility Seed Grant. Supported in part by the American Lung Association Innovation Award and National Institutes of Health grants (R21AG081715, R21AI173900, and R35GM151028) to A.F.C.

## Author Contribution

E.O. contributed to the experiments, data analysis, and writing of the paper. A.V. and F.G.R.M. contributed to the writing of the paper. A.F.C. supervised the project and wrote the paper.

